# Dissecting serotype-specific contributions to live oral cholera vaccine efficacy

**DOI:** 10.1101/2020.08.20.259119

**Authors:** Brandon Sit, Bolutife Fakoya, Ting Zhang, Gabriel Billings, Matthew K. Waldor

## Abstract

The O1 serogroup of *Vibrio cholerae* causes pandemic cholera and is divided into Ogawa and Inaba serotypes. The O-antigen is *V. cholerae’s* immunodominant antigen, and the two serotypes, which differ by the presence or absence of a terminally methylated O-antigen, likely influence development of immunity to cholera and oral cholera vaccines (OCVs). However, there is no consensus regarding the relative immunological potency of each serotype, in part because previous studies relied on genetically heterogenous strains. Here, we engineered matched serotype variants of a live OCV candidate, HaitiV, and used a germ-free mouse model to evaluate the immunogenicity and protective efficacy of each vaccine serotype. By combining vibriocidal antibody quantification with single and mixed strain infection assays, we found that all three HaitiV variants - Inaba^V^, Ogawa^V^, and Hiko^V^ (bivalent Inaba/Ogawa) - were immunogenic and protective, suggesting the impact of O1 serotype variation on OCV function may be minimal. The potency of OCVs was found to be challenge strain-dependent, emphasizing the importance of appropriate strain selection for cholera challenge studies. Our findings and experimental approaches will be valuable for guiding the development of live OCVs and oral vaccines for additional pathogens.

## Introduction

The human bacterial pathogen *Vibrio cholerae* causes cholera, a severe and potentially fatal diarrheal disease. In the small intestine, *V. cholerae* produces cholera toxin (Ctx), an AB_5_ toxin that induces ion imbalances and a secretory response that largely accounts for the massive fluid loss associated with cholera^1^. Although effectively treated with rehydration therapy, cholera remains a threat to public health. The disease is endemic in over 50 countries, and is especially dangerous where access to clean water and sanitation remains limited^1^. There are an estimated 3,000,000 cases and ~100,000 deaths due to cholera worldwide each year^2^. The magnitude of this threat has propelled interest in understanding *V. cholerae-*host immune system interactions for the refinement of oral cholera vaccines (OCVs), an important frontline intervention to reduce both cholera incidence and transmission^3^.

Both killed (inactivated) and live OCV formulations have been developed. Killed whole-cell OCVs (e.g. Shanchol), which consist of a mixture of heat or formalin-inactivated *V. cholerae* strains, have been generally efficacious in both endemic and epidemic settings^4^. However, inactivated vaccines have limited efficacy in young children (<5 years old), who are most susceptible to severe cholera, and require multi-dose immunization regimens for long-lived immunity, although recent studies suggest that single or higher dose schedules may still offer shorter term protection^5–8^. In contrast to killed OCVs, live OCVs are likely to be more effective after a single dose and in young children, since such vaccines more closely mimic authentic infection; during their *in vivo* replication in the intestine, live OCVs produce intact antigens, including infection-induced colonization factors, such as the toxin co-regulated pilus, that are targets for protective immunity^9–11^. However, to date, no live OCV is approved for use in cholera endemic countries, highlighting the need for development of new live OCVs for global public health. There is a live OCV (Vaxchora) that is commercially available in the USA, but its indication is limited to travel-related use^12^.

Although there are >200 known *V. cholerae* serogroups, all pandemic cholera has been caused by O1 serogroup *V. cholerae*^13^. Continued evolution of this dominant *V. cholerae* pandemic serogroup has given rise to several genetically and phenotypically distinct *V. cholerae* lineages (i.e. biotypes). Classical biotype *V. cholerae*, now thought to be extinct, likely caused the 1^st^-6^th^ cholera pandemics. The ongoing 7^th^ cholera pandemic, which began in 1961, is caused by the 7^th^ pandemic El Tor (7PET) biotype of O1 *V. cholerae*. Extensive genomic analyses of 7PET *V. cholerae* have demonstrated additional changes acquired by more recent clinical isolates, including the *ctxB7* allele of Ctx and the SXT antibiotic resistance element^14–16^. These “Wave 3” 7PET strains are now the primary cause of cholera worldwide, and have caused dramatic outbreaks in Haiti (2010-2019) and Yemen (2017-present)^17,18^.

The O-antigen moiety of lipopolysaccharide (LPS) is thought to be the primary antigenic determinant of protective immunity resulting either from O1 *V. cholerae* infection or vaccination^19^. The O1 serogroup includes two serotypes, Ogawa and Inaba, which differ by the presence or absence, respectively, of a methyl group on the terminal sugar of the LPS O-antigen (Figure 1A)^1,20–22^. Both Ogawa and Inaba strains cause epidemic cholera and circulate globally, often replacing each other in cyclical outbreaks in the same region^23–25^. Studies of natural infection indicate that immune responses and subsequent protection against future infection are strongest against the homologous (initial) serotype^26^. However, the relative potency of the cross-protectiveness of these responses, particularly whether one serotype confers greater cross-protectivity, remains unclear. Studies on this topic have not used matched isogenic strains to investigate the impact of variation in this immunodominant antigen in isolation from the complex suite of *V. cholerae* virulence and colonization factors. Similarly, while the serotype/biotype landscape of current OCVs is varied, ranging from monovalent (Vaxchora, classical Inaba) to trivalent (Shanchol, classical and El Tor Ogawa/Inaba/O139) formulations, evidence for their cross-serotype protective capacity from controlled comparisons of vaccines of varying serotypes is lacking. Understanding these concepts could guide design of a more potent O1 OCV against both serotypes.

**Figure 1.**
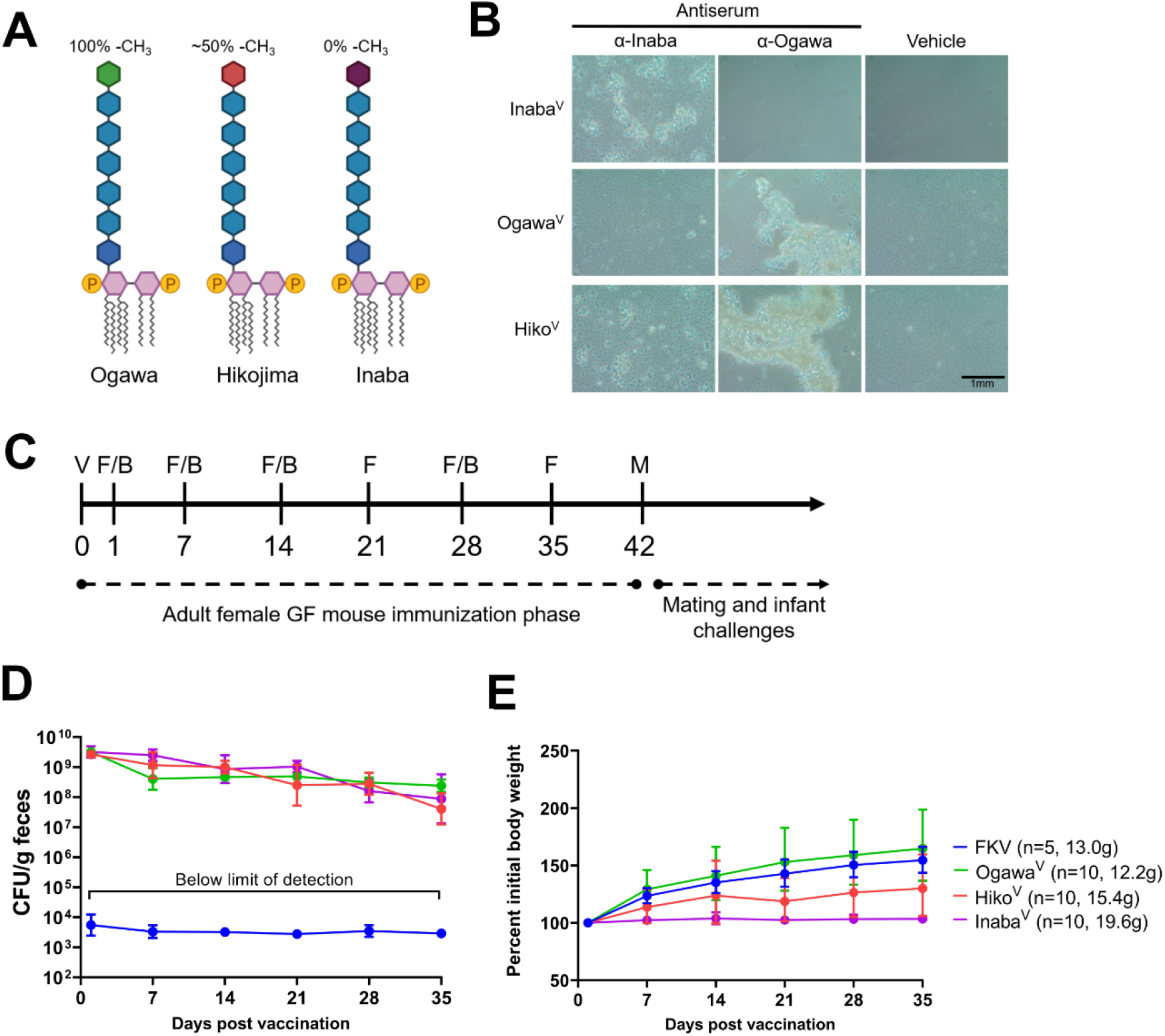
Vaccine shedding and bodyweights of adult germ-free mice orally immunized with isogenic vaccine serotype variants. (A): Schematic of the three known O1 *V. cholerae* serotypes. Internal and terminal perosamine residues are indicated by blue or otherwise colored diamonds, respectively, with the approximate degree of methylation of the terminal perosamine shown. (B): Representative slide agglutination of the three HaitiV vaccine serotypes. (C): Oral immunization and sampling regime for adult mouse phase of this study. V – oral vaccination, F – fecal pellet collection, B – blood sample collection, M – mating. (D) Fecal shedding and (E) bodyweight of mice orally immunized with a single dose of the indicated inactivated or live vaccine strain at Day 1.

O1 serotypes are determined by the activity of the O-antigen methyltransferase WbeT (previously known as RfbT), and loss-of-function mutations in this enzyme produce Inaba strains^27^. In rare cases where WbeT function is impaired, but not eliminated, Hikojima strains, which simultaneously produce methylated (Ogawa) and un-methylated (Inaba) LPS, can arise^27,28^. Although Hikojima is thought to be an unstable phenotype, Hikojima strains have historically and recently been isolated in the clinic^29–31^, and stable Hikojima-generating point mutations in WbeT have recently been identified^32,33^, raising the idea that an OCV bearing this bivalent O1 serotype could elicit superior cross-serotype protection while retaining the manufacturing advantages of a single strain vaccine. To this end, an inactivated Hikojima vaccine (Hillchol) has been produced, with promising early clinical results suggesting non-inferior immunogenicity compared to Shanchol^34^.

We recently described a new live-attenuated OCV candidate, HaitiV, an engineered derivative of a toxigenic Wave 3 7PET O1 Ogawa *V. cholerae* clinical isolate (HaitiWT) from the 2010 Haiti cholera epidemic. HaitiV contains a set of genetic modifications that reduce its potential reactogenicity and enhance its biosafety, and allow it to over-produce the non-toxic B subunit of Ctx to boost immunogenicity^35^. In two mouse models, we showed that HaitiV is immunogenic and elicits robust protective adaptive immune responses^36,37^. In addition to its function as a conventional live OCV, HaitiV also has an unprecedented, rapid-acting function that protects animals against lethal *V. cholerae* challenge within 24 hours post-immunization. Thus, HaitiV could be an OCV with both short- and long-term protective functions^35^. Here, to understand how vaccine serotype influences the generation of serotype-specific vibriocidal antibodies and protective immunity, we engineered isogenic Inaba, Ogawa and Hikojima variants of HaitiV. The protective capacities of these vaccines were tested in a germ-free (GF) mouse OCV immunization model using isogenic Inaba and Ogawa challenge strains. All three serotype vaccines functioned as excellent OCVs, with subtle but detectable differences in immunogenicity and protective efficacy. Our study provides insight into the *in vivo* biology of *V. cholerae* serotypes, demonstrates the utility of the GF mouse platform for rapidly assessing the OCV candidates, and offers guidance for design of future OCV trials.

## Results

### Generation of genetically matched HaitiV serotype variant strains

Among the suite of genetic modifications in HaitiV is a deletion of the recombinase *recA* (VC0543), which limits the vaccine’s capacity to acquire new genetic material, but is required for engineering mutations by homologous recombination. To further modify HaitiV, we restored a precursor of HaitiV to *recA*^+^. Subsequently, the hemolysin *hlyA* (VCA0219) was deleted, since HlyA is a suspected *V. cholerae* virulence factor^38^. We then introduced the following reported point mutants of WbeT (VC0255) at Ser158 in the original Ogawa variant of HaitiV: S158F (Hikojima), and S158P (Inaba)^32^. Finally, *recA* was deleted from each strain to yield three HaitiV-derived isogenic ∆*hlyA*/∆*recA* strains: Ogawa^V^ (WbeT^S158^), Hiko^V^ (WbeT^S158F^), and Inaba^V^ (WbeT^S158P^) (Figure 1A). Each vaccine strain’s serotype was confirmed by slide agglutination (Figure 1B).

### Vaccine serotype influences the specificity of vibriocidal responses

We used the GF adult mouse oral immunization model to compare the relative potency of the immune responses elicited by these three HaitiV serotype variants in adult female mice^36,39,40^. 3-6-week-old female GF mice (n=5/group) were immunized with a single oral dose of 10^9^ CFU of live Ogawa^V^, Inaba^V^, Hiko^V^ or formalin-inactivated Hiko^V^ (FKV) (Figure 1C). All three live variants were shed in feces (i.e. colonized) at equivalent levels with comparable kinetics over the course of the study (Figure 1D). Although there was some variance in mean initial weights of each group due to mouse availability, mice weights remained stable or increased over the course of the study, indicating that prolonged colonization by all 3 vaccine serotypes is safe (Figure 1E).

We next gauged immune responses to the vaccine variants by measuring vibriocidal antibody titers (VATs) in serum samples from the mice. VATs are a strong clinical correlate of protection in human *V. cholerae* infections and report on serotype-specific antibody responses against Ogawa and Inaba target strains^19^. All but two (28/30) mice immunized with live vaccines seroconverted (>4x from baseline VAT) against at least one O1 serotype within 14 days post-immunization, with the remaining two seroconverting by Day 28 (Figure 2). Anti-Ogawa responses were comparable between the three vaccine variants and ≥80% of animals responded to vaccination (Fig 2A). In contrast, anti-Inaba responses in Ogawa^V^-immunized mice were of lower titer than those in Inaba^V^- or Hiko^V^-immunized mice and were not detected in some animals 14 and 28 days post-vaccination (Fig 2B), suggesting that vaccine strain serotype biases the potency of serotype-specific vibriocidal immune responses. Only one FKV-immunized mouse seroconverted, demonstrating that the inactivated vaccine is markedly less immunogenic in this model. In a separate GF mouse cohort, we observed similar responses to *hlyA*+ Ogawa^V^ as the *hlyA* mutant, indicating that deletion of this locus did not broadly impact immunogenicity (Supplementary Figure 1).

**Figure 2.**
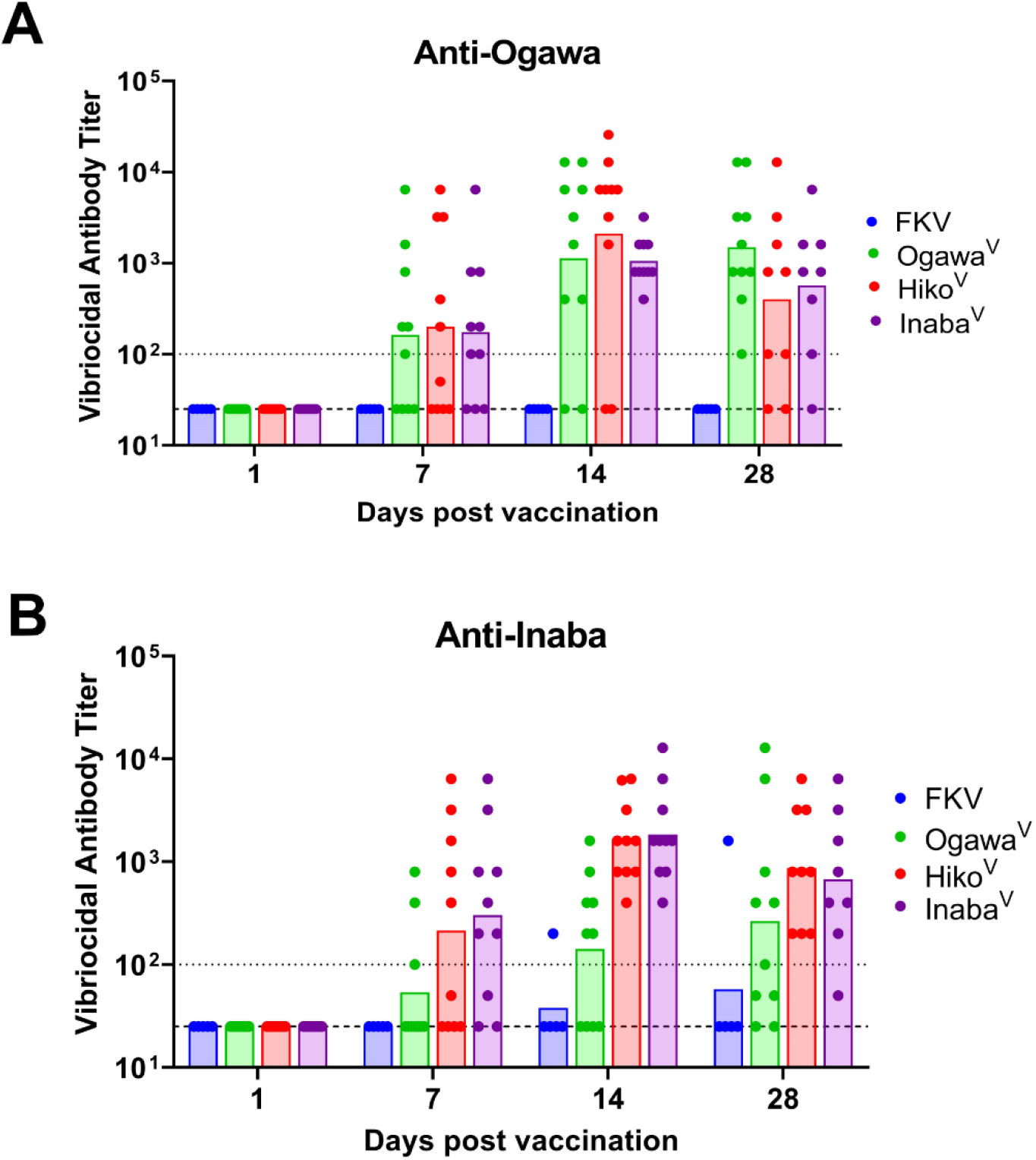
Vibriocidal antibody titers in adult germ-free mice orally immunized with isogenic vaccine serotype variants. Anti-Ogawa (A) and - Inaba (B) titers are plotted as geometric means of each group with individual values for each mouse shown. Individual values correspond to the highest dilution at which vibriocidal activity was observed. The lower dotted line represents the lower limit of detection (1:25 serum dilution), with the upper dotted line indicating the seroconversion threshold (4-fold increase over the baseline limit of detection).

### Vaccine serotype influences vaccine protective efficacy

VATs are not a direct measure of vaccine protective efficacy, and adult mice are refractory to cholera-like illness. To assess whether the immune responses in the vaccinated female adults were protective, and whether protection was biased by the serotype of the vaccine strain, we tested the susceptibility of their neonatal progeny to lethal challenge with either HaitiWT Ogawa (Ogawa^WT^) or Inaba (Inaba^WT^) isogenic challenge strains (Supplementary Figure 2). To control for litter-to-litter variations in maternal care and immune responses, which could affect comparisons, each litter was randomly split into two groups of pups, which received either a lethal Ogawa^WT^ or Inaba^WT^ challenge (Figure 3A). Pups in all three live vaccine groups were significantly protected from both death and diarrhea caused by challenge with either serotype compared with pups in the FKV group that exhibited similar kinetics of mortality as pups of unvaccinated dams, indicating that the presence of VATs (i.e. seroconversion) is tightly correlated with protection in this model (Figure 3BC)^36,37^. However, the protective efficacy of the three vaccine serotypes differed depending on the serotype of the challenge strain. All three live vaccines provided similar protection against Inaba^WT^ challenge (Figure 3B), but pups from Ogawa^V^-immunized dams were protected from death and diarrhea to a greater extent than pups from the other two groups against Ogawa^WT^ challenge (Figure 3C, p = 0.0033 and 0.0026 for diarrhea incidence in pups from Ogawa^V^ dams versus Hiko^V^ and Inaba^V^ dams, respectively). These findings suggest that Ogawa^V^ elicits more potent protective immune responses to homologous challenge and that vaccine serotype modifies the protective capacity of OCVs. In contrast to their differential protective capacities, all three vaccines conferred similar levels of colonization suppression (Fig 3BC), illustrating that suppression of colonization is not strictly equivalent to protection against disease.

**Figure 3.**
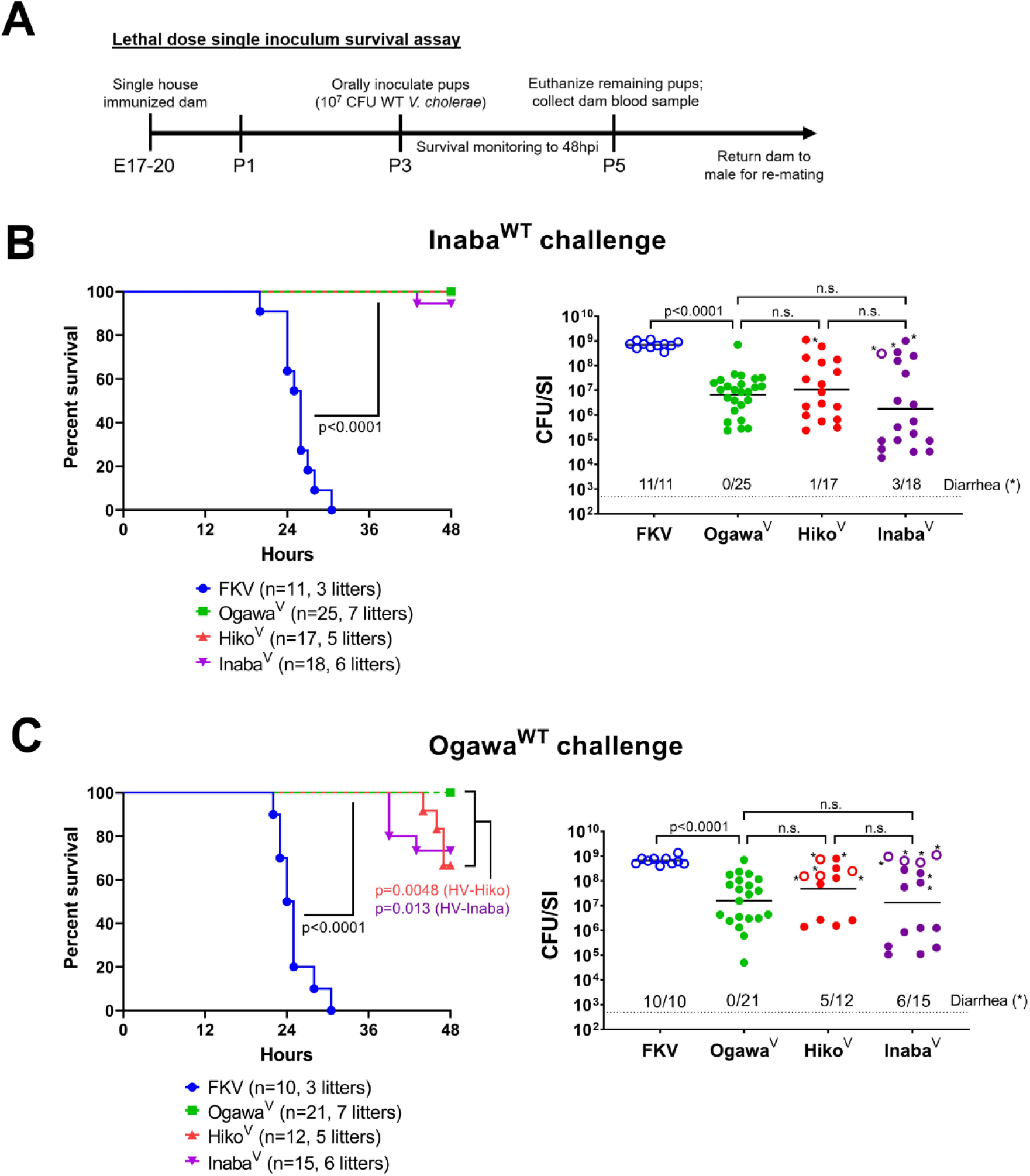
Protective efficacy of isogenic vaccine serotype variants using isogenic single strain challenges in pups from immunized dams. (A) Timeline of challenge experiments depicted in Figures 3, 4 and 6. For Figure 3 and 4, P3-4 pups in each litter were randomly assigned to receive a lethal dose of either Inaba^WT^ (B) or Ogawa^WT^ (C). Panels on the left depict survival kinetics for challenged pups from the indicated vaccine groups, with associated median survival times and sample sizes. P-values were determined by the Mantel-Cox test. Panels on the right show small intestinal (SI) *V. cholerae* burden in the same pups at the time of death (open circles) or at assay endpoint (48 hpi, closed circles). Burden is plotted as CFU/SI and pups with visible signs of diarrhea at the time of sacrifice are marked with an asterisk. P-values were determined by the Mann-Whitney U test.

We also analyzed the expansion of Inaba^WT^ or Ogawa^WT^ within each litter to counter potential confounding effects of variations in maternal care between litters. The *in vivo* expansion of the challenge strains was estimated by dividing the number of CFU recovered from the intestine by the CFU in the inocula, yielding a fold replication over inoculum (FROI) index. FROI comparisons revealed that litters from FKV, Ogawa^V^ and Hiko^V^ immunized dams did not display differential expansion of either the Inaba or Ogawa challenge strain (Figure 4A); in each group, litters showed comparable replication of either serotype. In contrast, in pups from all six litters from Inaba^V^ immunized dams, Ogawa^WT^ expanded to a greater extent than Inaba^WT^, suggesting that Inaba^V^-induced immune responses have diminished capacity to suppress the expansion of virulent Ogawa versus Inaba *V. cholerae* in the intestine. There was a statistically significant, but modest negative correlation between both peak or post-challenge mean dam VAT and FROI in the associated litters (Figure 4BC), suggesting that these metrics of vaccine potency are related in GF mice. Significant correlations between serotype-specific VAT and serotype-specific expansion were not detected in all groups, possibly due to insufficient statistical power (Supplementary Figure 3A-D). Importantly, other metrics such as dam bodyweight were not correlated to FROI, indicating the specificity of VAT correlations with pathogen replication in the GF mouse OCV model (Supplementary Figure 3EF).

**Figure 4.**
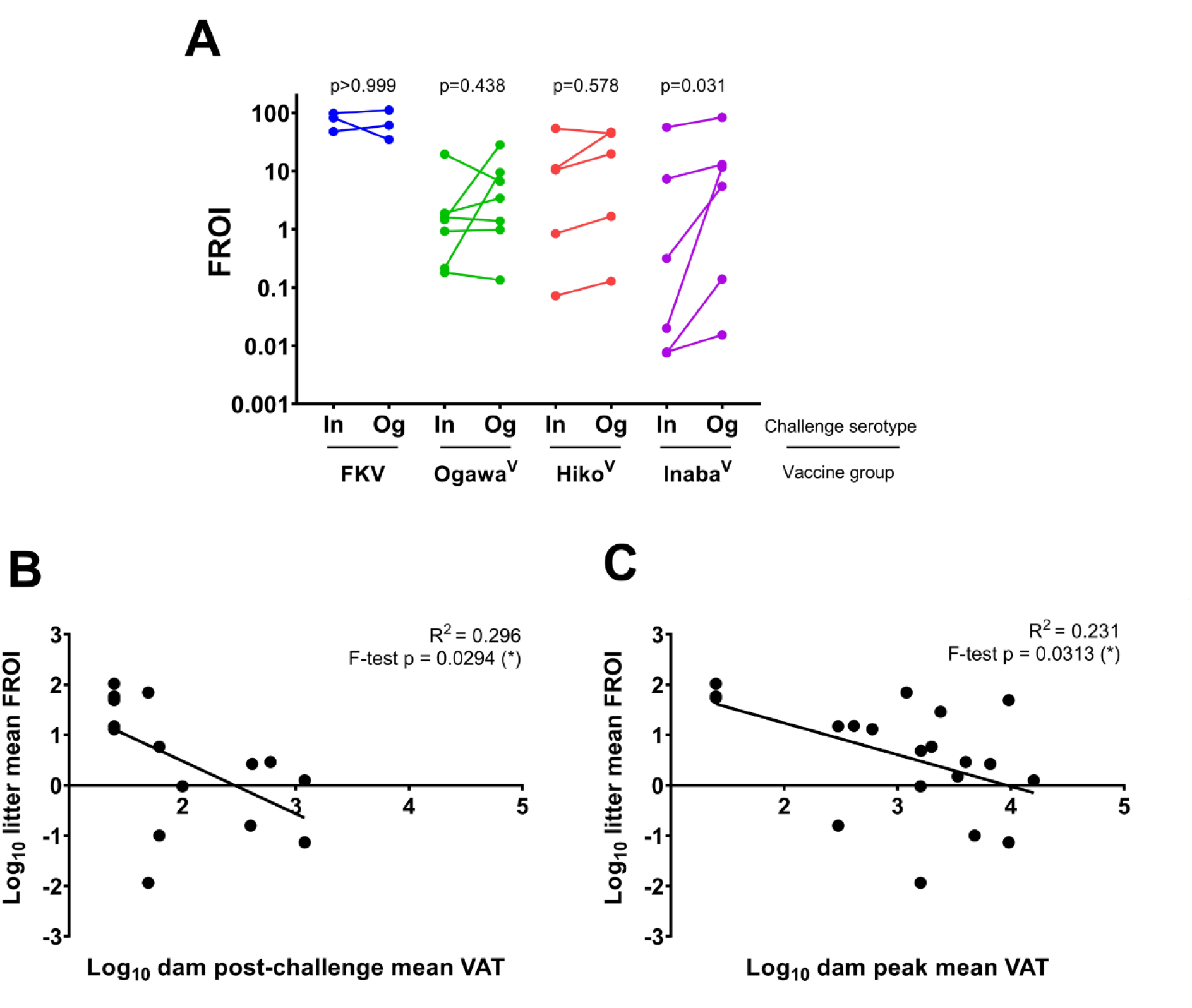
Replication of WT challenge strains in pups from dams immunized with isogenic serotype variants of OCV. (A): Fold replication over inoculum (FROI) for the indicated challenge strain (In: Inaba^WT^, Og: Ogawa^WT^) using colonization data from right panels in Figure 3. Solid lines connect FROI values for In- or Og-challenged pups in the same litter. P-values were calculated by the Wilcoxon matched-pairs signed rank test. (B): Correlation of litter-specific FROI with the post-challenge mean vibriocidal antibody titer (VAT) in the dam. Mean post-challenge VAT was determined by averaging the anti-Inaba and - Ogawa VAT values from the blood sample taken at the time the dam’s litter was euthanized. (C): Correlation of litter-specific FROI with the peak mean vibriocidal antibody titer (VAT) in the dam. Mean peak VAT was determined by averaging the highest measured anti-Inaba and - Ogawa VAT values from the mouse at any point during the study. The p-value of the fitted linear regression was calculated with the F-test.

To further probe the serotype-specific protective efficacy of a bivalent vaccine strain such as Hiko^V^, we carried out within-pup comparisons using competitive infections, thus controlling for pup-to-pup variations in maternal care. These experiments used a lower challenge inoculum (10^5^ CFU) composed of a 1:1 mixture of Ogawa^WT^ and Inaba^WT^ to robustly assay colonization in the absence of signs of disease (Figure 5A). Consistent with the single infection data, the total CFU burden of WT *V. cholerae* in pups from Hiko^V^ dams was significantly reduced (~100x) compared to pups from unimmunized dams (Figure 5B). Unexpectedly, in pups from unimmunized dams, the baseline competitive index (CI) was not 1, suggesting that Ogawa^WT^ has a modest competitive advantage in the infant mouse SI relative to Inaba^WT^ (Figure 5C). CIs in pups from Hiko^V^-immunized dams were significantly higher than in the controls, suggesting that Hiko^V^ vaccination elicits immune responses that are less potent at impeding Ogawa^WT^ expansion and/or more potent at impairing Inaba^WT^ replication (Figure 5C). These observations provide additional evidence that OCV serotype exerts serotype-specific impacts on *V. cholerae* fitness, even within a single animal.

**Figure 5.**
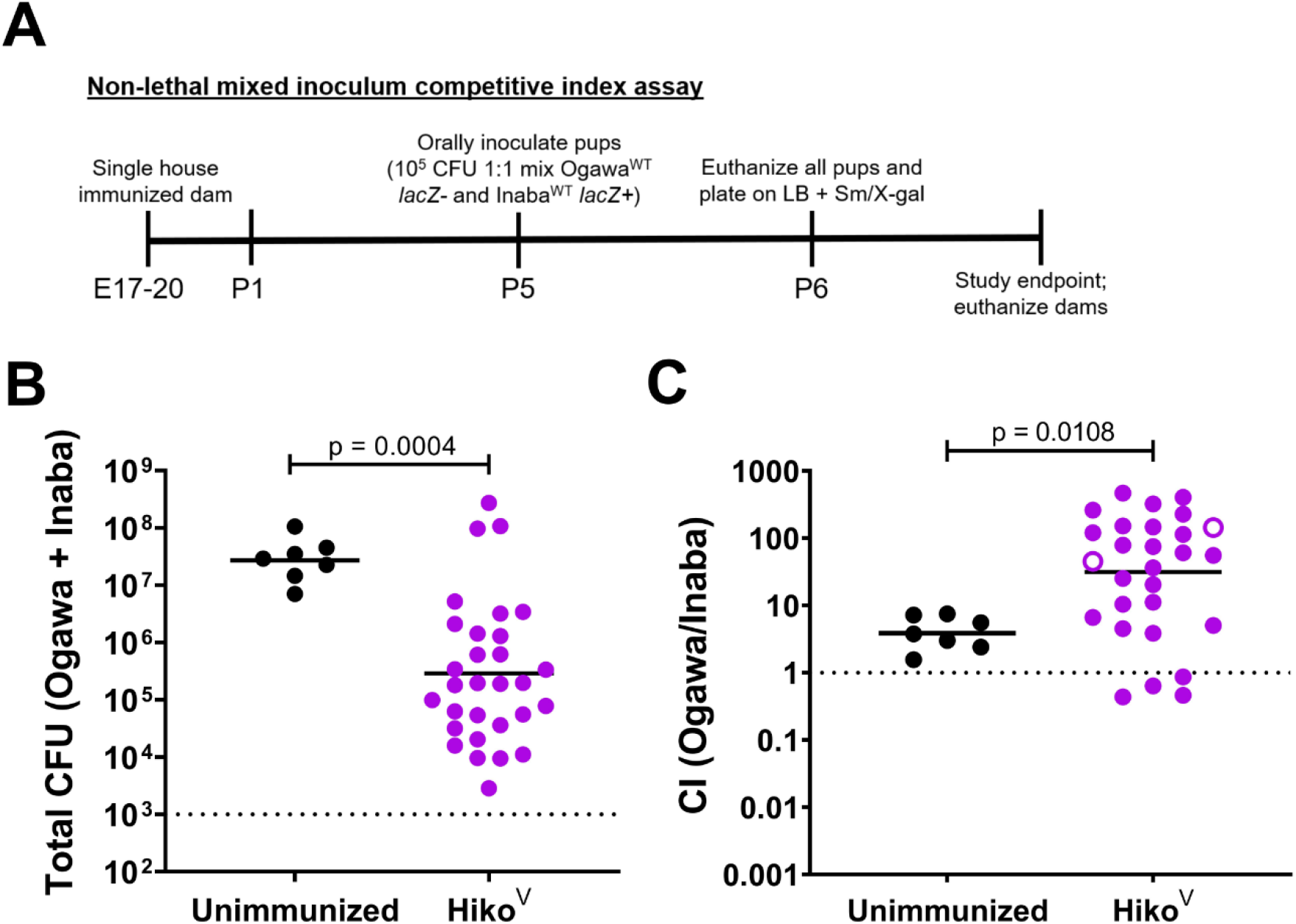
Mixed-strain challenges in pups from dams immunized with Hiko^V^. (A): Timeline of mixed strain challenge experiments. (B): Colonization burdens in P6 pups from unimmunized and Hiko^V^ vaccinated adults challenged with a 1:1 mixture of Inaba^WT^ and Ogawa^WT^. The dotted line indicates the limit of detection. (C): Competitive index (CI) values from pups in Panel B. Open circles denote pups where the denominator was 0 in the CI calculation, forcing an imputed 1 and hence representing the upper limit of detection. The dotted line indicates the line of parity (CI = 1). Pups with a total colonization burden of less than 5×10^3^ CFU in Panel B were excluded from Panel C. P-values were determined by the Mann-Whitney U test.

### Serotype-independent challenge strain properties impact the protective efficacy of Hiko^V^

To further explore the protective scope of the bivalent Hiko^V^ strain against diverse *V. cholerae* challenges, we immunized an additional cohort of GF mice with formalin-inactivated or live Hiko^V^. Hiko^V^ stably colonized the mice and the vaccinated animals gained weight and developed similar serum VATs as described above (Supplementary Figure 4). We challenged litters from these vaccinated dams with 7PET clinical strains from the last several decades to capture the evolutionary spectrum of the 7^th^ cholera pandemic. These included N16961, an Inaba Bangladeshi isolate from 1971^41^, PIC018, an Inaba Bangladeshi isolate from 2007^42^ along with the HaitiWT (isolated in 2010) derived strains Ogawa^WT^ and Inaba^WT^, used above (Supplementary Figure 2). Consistent with data from the previous cohort (Figure 3), pups from FKV-vaccinated dams were not protected against disease or colonization (Figure 6). In contrast, pups from live Hiko^V^-immunized dams were completely protected against the three contemporary isolates (PIC018, Ogawa^WT^ and Inaba^WT^) (Figure 6A). None of these animals developed diarrhea or died. Despite the equivalent clinical protection from these three challenge strains, Hiko^V^ vaccination suppressed colonization by PIC018, an Inaba strain, more potently than either Haitian serotype (Figure 6B). Conversely, pups from Hiko^V^-immunized dams displayed significantly lower levels of clinical protection (7/17 with diarrhea/died, p=0.0087 vs. Inaba^WT^) and colonization suppression when challenged with the early 7PET strain N16961 (Figure 6). These findings strongly suggest that strain-specific factors in addition to serotype-specific immune responses determine the protective efficacy of OCVs.

**Figure 6.**
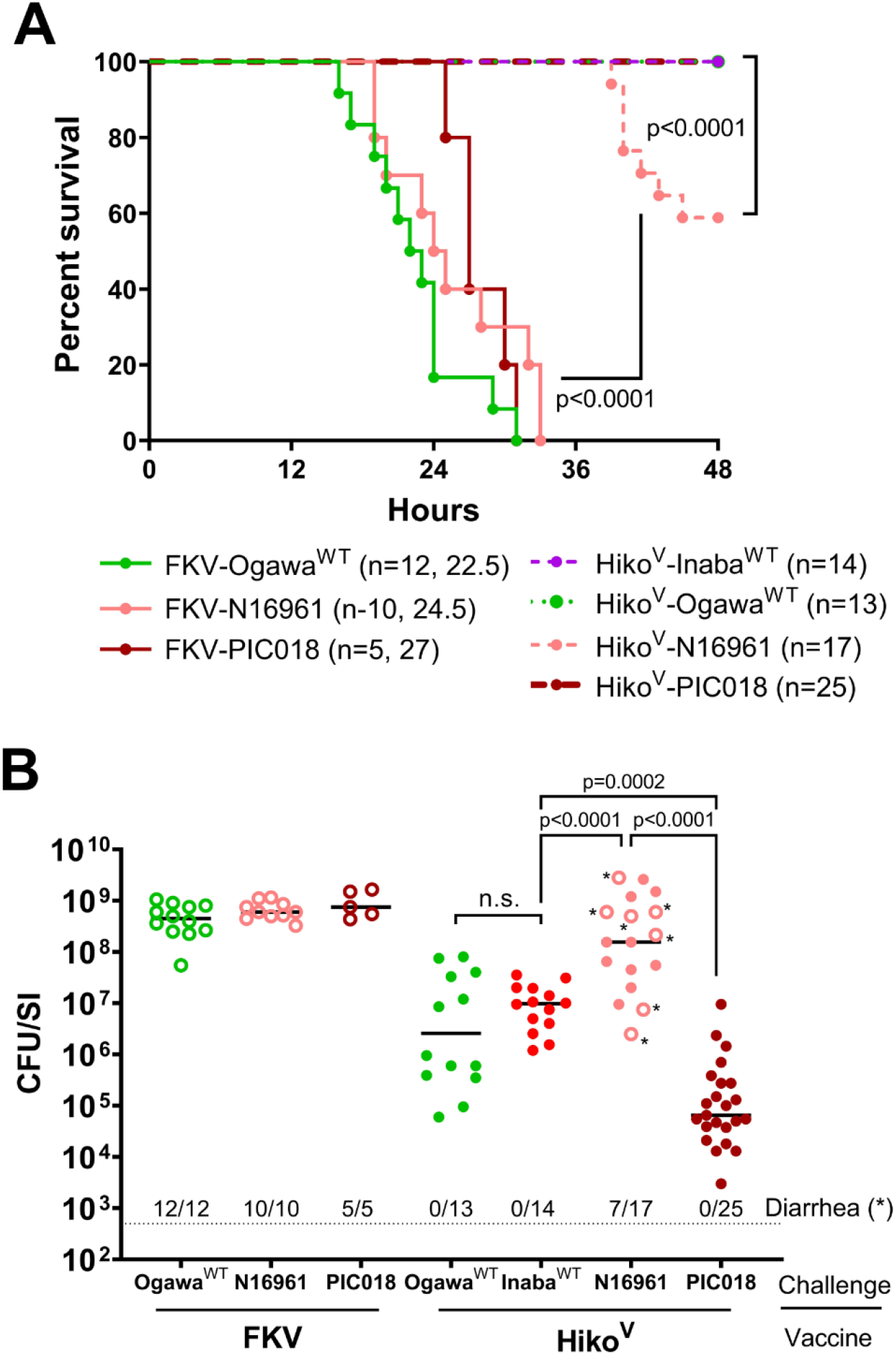
Diverse single strain challenges in pups from dams immunized with Hiko^V^. (A) Survival kinetics in P3-4 pups from dams immunized with formalin-killed (FKV) or live Hiko^V^ challenged with the indicated WT *V. cholerae* strain. The sample size and median survival times, where calculable, are indicated. (B) Intestinal *V. cholerae* burden in the pups from Panel A at the time of death (open circles) or at assay endpoint (48 hpi, closed circles). Burden is plotted as CFU/SI and pups with visible signs of diarrhea at the time of sacrifice are marked with an asterisk. P-values were determined by the Mann-Whitney U test.

## Discussion

The Inaba and Ogawa serotypes of O1 serogroup *V. cholerae*, which were initially described over a century ago, continue to cause virtually all pandemic cholera^43^. Based on knowledge that the O1 O-antigen is a critical target of protective immunity against cholera and that the methylation that distinguishes Ogawa from Inaba strains can impact anti-*V. cholerae* immune responses, we engineered genetically matched serotype variants of a live OCV candidate, HaitiV, as well as isogenic Ogawa and Inaba WT challenge strains, to determine which, if any, O1 serotype would be the most immunogenic and protective. We hypothesized that the bivalent Hikojima serotype would be the most effective OCV formulation, as it presents both Inaba and Ogawa antigens. We found that all three HaitiV variants - Inaba^V^, Ogawa^V^, and Hiko^V^ - were both immunogenic and protective in the GF mouse model. Given that we observed relatively minor differences between the vaccines, we suggest that the impact of O1 serotype variation in OCV design may be minimal, and our data did not consistently identify a superior serotype across the assays we performed. However, a striking result from this study was that all three live vaccines were far more immunogenic than a formalin killed version of Hiko^V^, strongly supporting the idea that a live OCV has great potential for control of cholera.

In the challenge assays, all three live vaccines protected nearly all pups from Inaba^WT^ challenge, but Ogawa^V^ was superior to either Inaba^V^ or Hiko^V^ in the animals challenged with Ogawa^WT^, even though the latter two vaccines elicited nearly equal anti-Ogawa VATs as Ogawa^V^. Despite protecting as well as Inaba^V^ and Hiko^V^ against Inaba^WT^ challenge, Ogawa^V^ induced lower anti-Inaba VATs than the other two vaccines, underscoring the complexities of experimental markers of protective immunity and suggesting that finer-scale metrics to evaluate vaccine efficacy would be valuable. This is especially important since immune and protection metrics both report on vaccine potency, but the translational insight imparted by these assays in relation to each other is not entirely clear. Given the tight range of protective efficacy of all three vaccines (85-100%), it was not possible to correlate VAT level and protection in the animals in our studies, as had been done in humans, beyond the observation that seroconversion was tightly associated with protection^44,45^. The split-litter challenges showed that a surrogate measure of efficacy, the FROI bacterial replication value, correlated with the level of circulating vibriocidal antibodies in the dam. This is consistent with knowledge that immunity to cholera is primarily driven by anti-bacterial effects^46^. Similarly, our use of mixed isogenic challenge strains allowed us to directly measure the relative replication (CI) of Inaba and Ogawa *V. cholerae* strains against each other in the same intestinal environment. This experimental format revealed that immunization with Hiko^V^ elicited immune responses that skewed the relative expansion of the Ogawa and Inaba challenge strains. In conjunction with the observation of serotype-bias in FROI, these data support the idea that anti-O-antigen antibodies are the direct effectors responsible for vaccine-mediated suppression of colonization^5,47^ and suggest that determination of the molecular bases of serotype-biased V. cholerae intestinal replication is warranted.

An important caveat underlying our findings is that GF mice lack commensal gut microbes and could have altered immune responses to immunization, especially considering recent studies that have highlighted how *V. cholerae*-microbiota interactions can influence colonization, disease, and development of anti-*V. cholerae* immunity^48–52^. However, the strong association of VAT induction with protection and superior efficacy of live over inactivated vaccine strains in this model, and our similar findings of OCV function in mice with transiently disrupted microbiomes reinforces the idea that the GF mouse live OCV model holds substantial translational promise^36,37^.

Investigations of the influence of the O1 serotypes on natural *V. cholerae* infection or OCV function in the literature are sparse, and sometimes conflicting. Surveillance-based natural infection studies conducted in Bangladesh, where cholera is endemic, have suggested that Inaba cholera infections confer stronger protection against future exposure to Ogawa *V. cholerae* than the reverse heterologous re-infection^25,26^. Conversely, an early volunteer challenge study with WT *V. cholerae* suggested that Ogawa-stimulated immunity against both homologous and heterologous serotype re-challenge was non-inferior to that conferred by Inaba^41^. In addition, recent analyses of immune responses during cholera suggest that Ogawa infections may elicit stronger cross-serotype reactive immune responses than Inaba infections^53^. There are several reasons that may explain why these studies have yielded disparate conclusions. First, the relatively small numbers of re-infected subjects in both challenge and surveillance studies limit their robustness. Additionally, serotype dynamics during cholera epidemics can range from clonal domination to co-circulation of Inaba and Ogawa strains, with local outbreaks often “switching” from Ogawa to Inaba and back again, complicating the interpretation of serotype-specific disease incidence in surveillance studies by impacting serotype-specific re-infection frequencies^24,25,54–57^. This is further confounded by the knowledge that serotype switching can be driven by selective pressure from serotype-specific antibodies, sometimes within the same host, making it likely that changes in population-level serotype-specific immunity drive serotype switching during outbreaks^29,58,59^. Lastly, no study has directly compared the immunogenicity or protective efficacy of Inaba and Ogawa OCVs with otherwise identical genetic backgrounds, a crucial control in the context of a continuously evolving pathogen such as *V. cholerae*. Our findings thus augment epidemiological observations and provide a needed framework to benchmark the performance of all three known O1 *V. cholerae* serotypes as OCVs.

Since we cannot predict *a priori* which serotype will cause an outbreak, and given that the reports described above conflict on whether Ogawa or Inaba vaccines are preferable, our data suggest that the development of a live Hikojima OCV would be the most risk-averse approach to next-generation OCV design. Hiko^V^ was largely non-inferior to Inaba^V^ and Ogawa^V^ in our study, presents both Ogawa and Inaba antigens, and, as a single strain formulation, could simplify the OCV manufacturing process. It will be of interest to investigate how different Hikojima-generating alleles of *wbeT* that give rise to skewed (not 1:1) Inaba:Ogawa O-antigen ratios influence serotype switching, immunogenicity and OCV efficacy.

Finally, the controls and experimental schemes we used should be valuable for the design of future investigations of OCVs, not only in GF mice, but potentially also in humans, as cholera is one of the few infectious diseases for which there is a human challenge model^60^. For example, using mixed-challenge inocula (e.g. Inaba and Ogawa *V. cholerae*) could enable assessment of the within-host relative fitness of the two major *V. cholerae* serotypes in immunized human volunteers. Our observation that the apparent potency of OCVs is challenge strain-dependent (Figure 6) also emphasizes the need for careful selection of relevant strains in animal and human investigations of OCVs. Similar considerations were taken when the early 7PET isolate N16961 was introduced as an updated challenge in 1980^41^. This isolate has since been used as the predominant human challenge strain, including the most recently reported volunteer challenge OCV trial^45^. Recent 7PET evolution over the last three decades has led to strains with altered virulence traits, and potentially immunogenicity^61–64^. Our data suggest that currently circulating late 7PET *V. cholerae*, which have never been used as challenge strains, may interact differently with OCV-induced immunity than early 7PET and 6^th^ pandemic isolates, and thus should be considered for use not only as next-generation live OCVs, but as challenge strains in future human volunteer studies. This change may advance the development of efficacious OCV candidates as well as yield mechanistic insights into immune responses to contemporary pandemic *V. cholerae.*

## Methods

### Bacterial strains and growth conditions

Bacteria were grown in lysogeny broth (LB) supplemented with the indicated antibiotics/compounds at the following concentrations: streptomycin (Sm, 200μg/mL), kanamycin (200μg/mL), carbenicillin (Cb, 50μg/mL), chloramphenicol (Cm, 0.75μg/mL) sulfamethoxazole/trimethoprim (SXT, 80 and 16μg/mL) and 5-bromo-4-chloro-3-indolyl-β-d-galactopyranoside (X-gal, 60μg/mL). For growth on plates, LB + 1.5% agar was used. Unless otherwise noted, strains were grown in liquid media at 37°C shaking at 200rpm. All *V. cholerae* strains in this study were spontaneous SmR derivatives of the wild-type. Bacterial stocks were stored at −80°C in LB with 35% glycerol. Strains and plasmids used in this study are listed in Supplementary Table 1 and 2, respectively.

### Serotype agglutination assay

To serotype vaccine and WT strains, triplicate 10μL drops of saturated overnight single colony cultures of each strain were spotted onto a glass slide and mixed with 5μL of anti-Inaba or anti-Ogawa sera (BD Difco) or a vehicle control (0.85% NaCl). Drops were then mixed with a sterile pipette tip and gently rocked for 30 seconds. Anti-Inaba serum strongly agglutinates Inaba and Hikojima *V*. *cholerae*. Anti-Ogawa serum strongly agglutinates Ogawa and Hikojima *V. cholerae*. Bacteria were imaged with a Nikon Eclipse TS100 inverted microscope at 100x magnification.

### Engineering HaitiV and HaitiWT derivatives

HaitiV and HaitiWT variants were created by conventional allelic exchange techniques as previously described^35^. Briefly, the HaitiV or HaitiWT *V. cholerae* derivative of interest was conjugated with SM10λpir *E. coli* bearing the suicide plasmid pCVD442 or pDS132 carrying the allele to be exchanged as well as 700bp of up- and downstream homology to the targeted genomic region. Conjugations were performed at a 1:1 donor:recipient ratio for 4 hours at 37°C and single crossovers were isolated by plating reactions on LB + Sm/Cb (pCVD442) or LB + Cm (pDS132) agar plates. To select for double crossovers, single crossover colonies were re-streaked on LB + 10% overnight at 30°C or were grown in LB + Sm/Cb for 4 hours at 37°C and then sub-cultured 1:100 statically in LB + 10% sucrose overnight at room temperature, followed by plating on LB + Sm. Colonies were then checked for Cb resistance by duplicate patching. Sm^R^/Cb^S^ colonies were screened by colony PCR (for *hlyA, lacZ* and *recA*) or Sanger sequencing (for *wbeT*) to identify colonies with the correct double crossover.

### GF mouse oral immunization scheme and sample collection

3-6-week old GF female C57BL/6 mice were obtained from the Massachusetts Host-Microbiome Center and housed in autoclaved cages with food and water given *ad libitum* in a non-gnotobiotic BL-2 facility on a 12-hour light/dark cycle for the duration of the study. On Day 0, mice were anesthetized with isoflurane and orally gavaged with 10^9^ CFU of overnight culture of one of the three serotype variants of live or formalin-killed (inactivated, FKV) HaitiV in 100uL 2.5% NaHCO_3_. Formalin inactivation was performed as previously described^35^. Immunized mice were monitored daily and weighed weekly. At the indicated timepoints in Figure 1A, a blood sample was collected by submandibular puncture for immunological assays and a fresh fecal pellet was collected from each mouse for plating on LB + Sm to enumerate HaitiV shedding. Blood samples were clotted at room temperature for 1 hour, centrifuged at 20000 x g for 5 minutes and supernatant (serum) stored at −20°C for subsequent analyses. Mice were co-housed according to vaccine group until Day 42, when mice designated for mating were re-housed with age-matched GF C57BL/6 males to initiate the mating and infant challenge study phase.

### Quantification of vibriocidal antibody titers

A complement-mediated cell lysis assay was performed to quantify vibriocidal responses in serum samples as previously described^40^. The clinical isolates PIC018 and PIC158 were used as the Inaba or Ogawa *V. cholerae* target, respectively. Seroconversion was defined as ≥4x increase in titer relative to the first measurement. A characterized mouse monoclonal antibody targeting *V. cholerae* O1 O-specific polysaccharide was used as a positive control for the vibriocidal assay^36^. Titers are reported as the dilution of serum causing a 50% reduction in target optical density compared to control wells with no serum added.

### Infant mouse single strain lethal challenge assay

Lethal dose single strain challenge assays were performed as previously described^36^. Pregnant dams were singly housed at E17-20 for delivery. At P3 (third day of life), pups were orally inoculated with 10^7^ CFU of the indicated WT *V. cholerae* strain in 50μL LB and returned to their dam. In split litters receiving Inaba^WT^ or Ogawa^WT^, pups were randomly assigned to each inoculum. Infected pups were monitored every 4-6 hours for onset of diarrhea and reduced body temperature. Once signs of disease were observed, monitoring was increased to 30-minute intervals until moribundity was reached, at which point pups were removed from the nest and euthanized for dissection, homogenization and plating of the small intestine (SI) on LB + Sm/X-gal for CFU enumeration. Pups that were alive at 48 hours post inoculation (hpi) were deemed protected from the challenge. Upon removal of the final pup in each litter, a submandibular blood sample was collected from the dam for vibriocidal antibody titer quantification. Fold replication over inoculum (FROI) values were calculated by dividing the total *V. cholerae* SI CFU burden at time of sacrifice by the inoculum. Mean FROI values were calculated by averaging FROIs from both surviving and succumbed pups in a given subgroup (i.e. Inaba or Ogawa-inoculated pups in a given litter or all pups in the litter). We excluded pups rejected by their dams from analyses due to our inability to attribute mortality to infection alone.

### Infant mouse mixed strain non-lethal competitive index assay

For non-lethal, competitive index (CI) infections, P5 (fifth day of life) pups were separated from their dams and orally inoculated with a 1:1 mix (total 10^5^ CFU) of HaitiWT Ogawa *lacZ-* and HaitiWT Inaba *lacZ+ V. cholerae*, a dose insufficient to cause disease or mortality but sufficient for robust intestinal colonization^65^. At 20 hpi, pups were euthanized for dissection and CFU plating of the SI on LB+Sm/X-gal for blue/white colony counting. CIs were obtained by dividing the ratio of white:blue (Ogawa:Inaba) colonies in the SI to the ratio of white:blue colonies in the inoculum.

### Statistical analysis

Statistical analyses were performed with Prism 8 (Graphpad). Survival curves were analyzed with the log-rank (Mantel-Cox) test and CFU burdens and CIs were compared with the Mann Whitney U test. Correlations between fold replication and VATs were generated by linear regression and statistically tested with the F-test. Bacterial replication within the same litter (FROI) was analyzed with the Wilcoxon signed-rank matched pair test. Differential incidence of diarrhea in challenged pups was analyzed with two-tailed Fisher’s exact tests. A p-value <0.05 was considered statistically significant.

### Animal use statement

This study was performed in accordance with the NIH Guide for Use and Care of Laboratory animals and was approved by the Brigham and Women’s Hospital IACUC (Protocol 2016N000416). Infant (P14 or younger) mice were euthanized by isoflurane inhalation followed by decapitation. Adult mice were euthanized at the end of the study by isoflurane inhalation followed by cervical dislocation.

## Acknowledgements

We thank members of the Massachusetts Host-Microbiome Center, especially Vladimir Yelisheyev and Rosa Perez Gonzalez, for assistance with GF mouse procurement and husbandry. We also thank the members of the Waldor lab for helpful discussions during this study and comments on the manuscript. Research in the Waldor Lab is funded by the National Institutes of Health (R01-AI-042347), the Wellcome Trust (218443/Z/19/Z) and the Howard Hughes Medical Institute. BS was supported by a fellowship from the Natural Sciences and Engineering Research Council of Canada (NSERC PGSD3-487259-2016).

## Author Contributions

Conceptualization: BS, BF, MKW. Methodology: BS, BF, MKW. Investigation: BS, BF, TZ, GB. Supervision: MKW. Visualization: BS. Writing (Original Draft Preparation): BS, BF, MKW. Writing (Review and Editing): BS, BF, TZ, GB, MKW.

