## Supplementary Information for "Dissecting serotype-specific contributions to live oral cholera vaccine efficacy"

**Supplementary Figures**

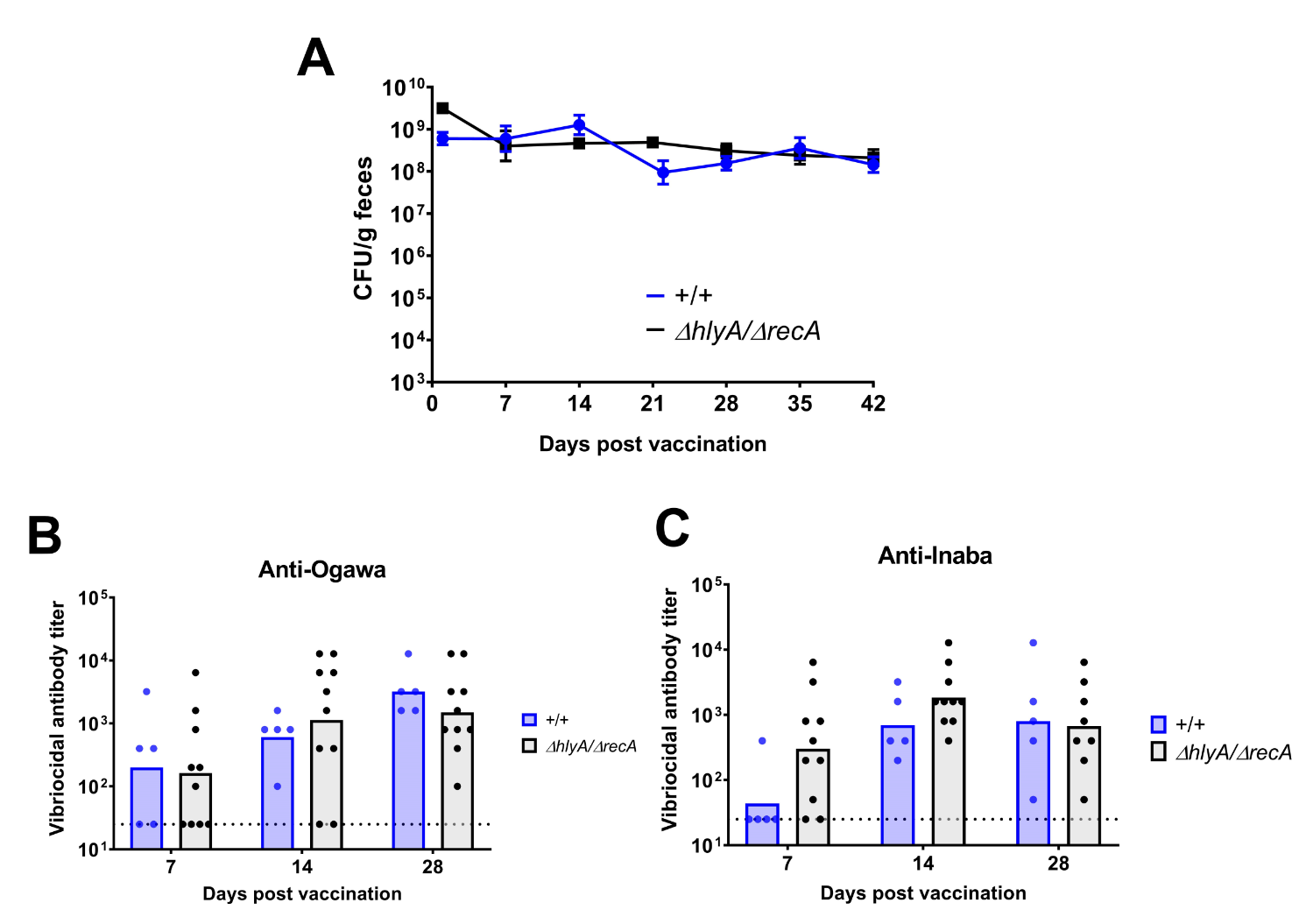

**Supplementary Figure 1. Effect of *hlyA* deletion on immunogenicity of HaitiV in adult germ-free mice.** (A): Fecal shedding of *hlyA+/recA+* (+/+) or Δ*hlyA/*Δ*recA* Ogawa HaitiV. (B) Anti-Ogawa and (C) anti-Inaba vibriocidal antibody titers in serum samples from the indicated timepoints in each vaccine group. Data for the Δ*hlyA/*Δ*recA* Ogawa^V^ mice is replicated from Figure 1 and 2 in the main text to allow comparison with mice immunized with the +/+ Ogawa^V^ strain.

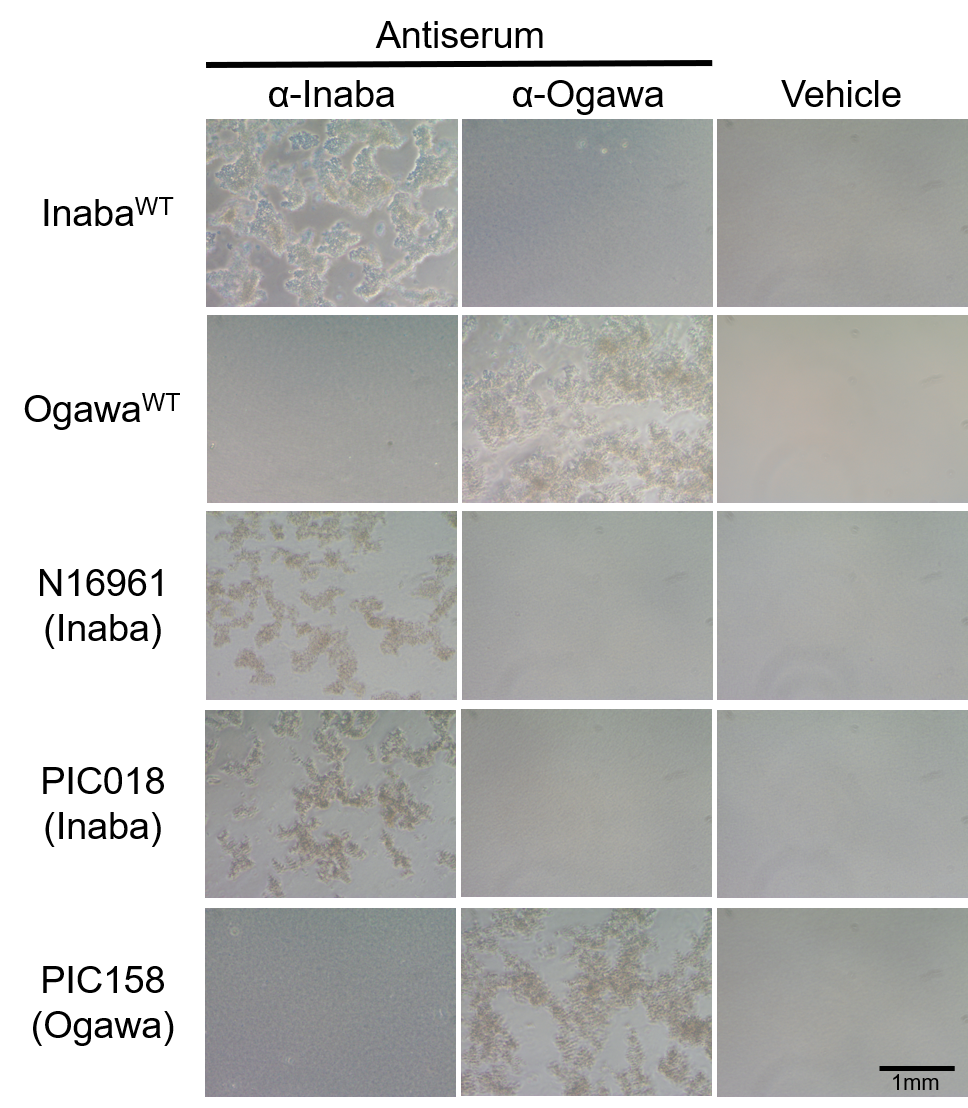

**Supplementary Figure 2. Slide agglutination of wild-type *V. cholerae* strains used in this study.** Representative images are shown.

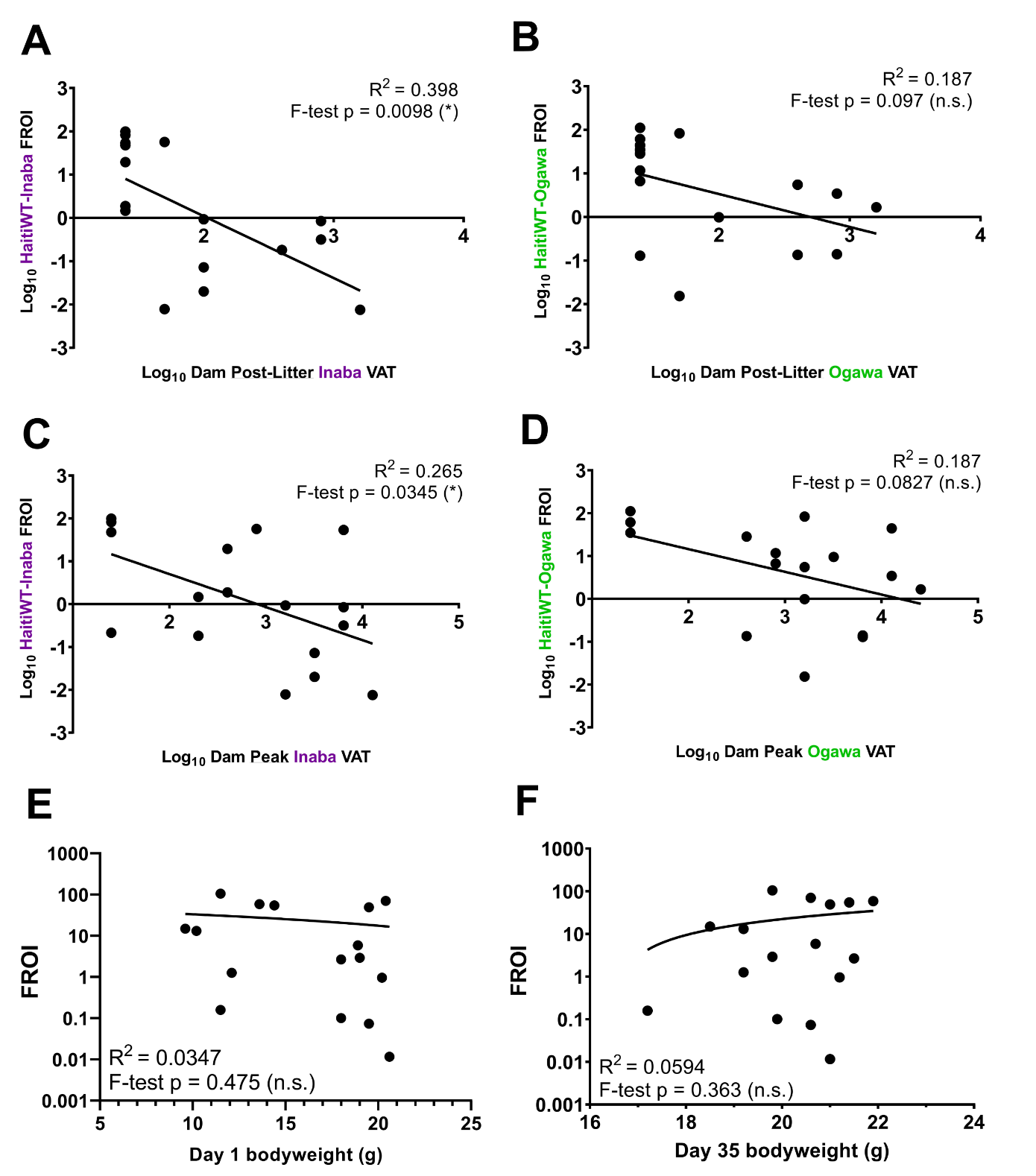

**Supplementary Figure 3. Correlation of fold replication values with dam anti-*V. cholerae* antibody responses or bodyweight.** Analysis was performed as in Figure 4BC in the main text, but without combining Inaba and Ogawa challenge FROI and VAT data. Shown are matched serotype FROI and VAT analysis for post-litter (A and B) or peak (C and D) VATs. To test the specificity of these relationships, mean litter FROI (combined across serotypes) was plotted against the assumed unrelated metric of dam bodyweight at Day 1 (E) or Day 35 (F) post-immunization.

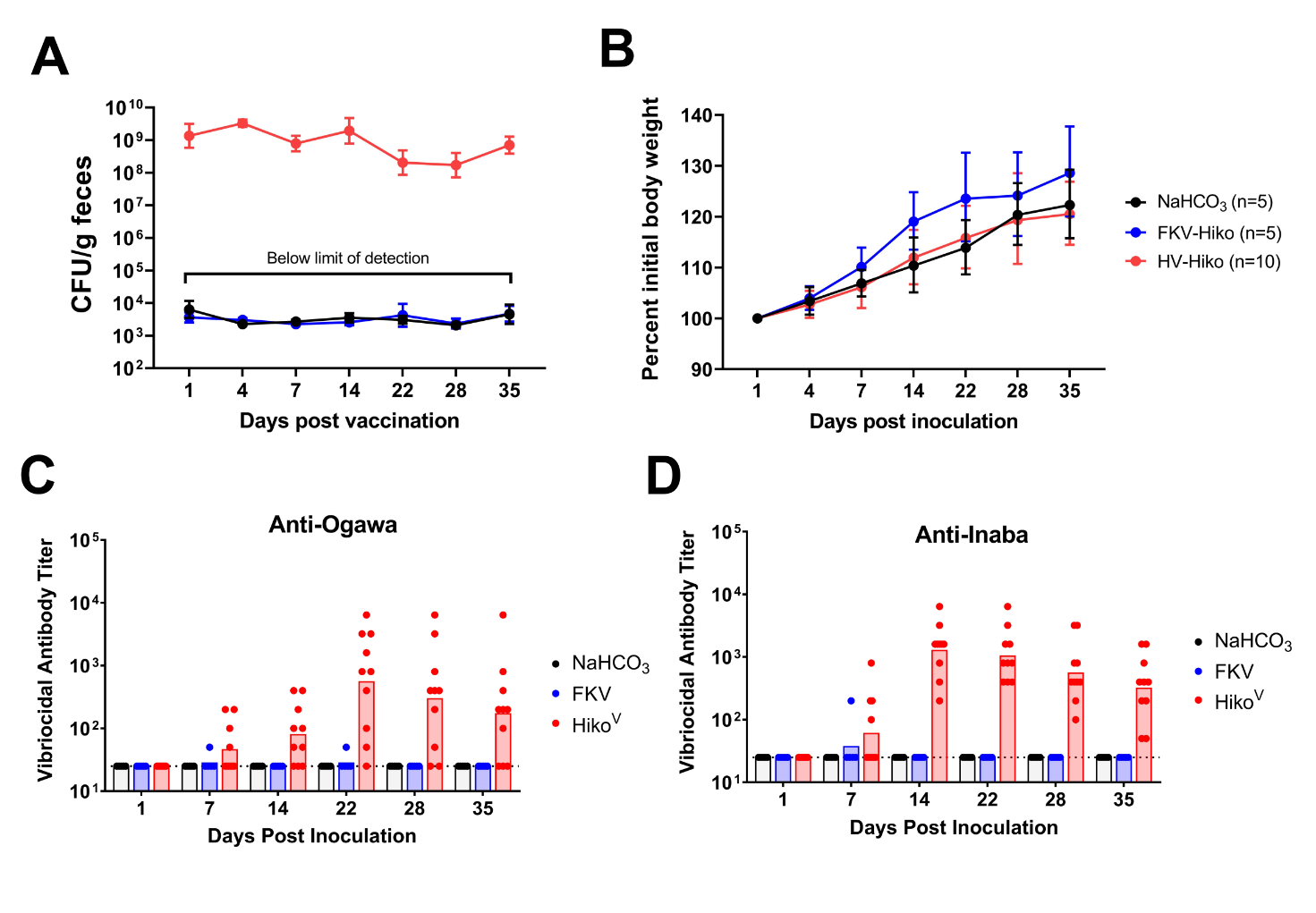

**Supplementary Figure 4. Overview of adult germ-free mice immunized with formalin-inactivated or live HaitiV-Hikojima.** Mice were immunized with buffer only (NaHCO_3_), inactivated (FKV) or live (Hiko^V^) vaccine at Day 0. Fecal shedding (A), bodyweight (B) and anti-Ogawa (C) and anti-Inaba (D) vibriocidal antibody titers were measured according to a similar sampling scheme to that described in Figure 1C in the main text.

**Supplementary Tables**

**Supplementary Table 1. Bacterial strains used in this study.** ET = El Tor, Sm = streptomycin, SXT = sulfamethoxazole/trimethoprim, R = resistant, S = sensitive.

| **Strain** | **Notes** | **Source** |
| --- | --- | --- |
| *E. coli* SM10pir | Donor strain for allelic exchanges | ^1^ |
| *V. cholerae* Ogawa^WT^ | HaitiWT, O1 Ogawa ET, SmR SXTR, *lac+ wbeT^S158^* | ^2^ |
| *V. cholerae* Ogawa^WT^ Δ*lacZ* | HaitiWT, O1 Ogawa ET, SmR SXTR, *lac-* (Δ*lacZ*) *wbeT^S158^* | This study |
| *V. cholerae* Inaba^WT^ | HaitiWT, O1 Inaba ET, SmR SXTR, *lac+ wbeT^S158P^* | This study |
| *V. cholerae* Ogawa^V^ +/+ | HaitiV, O1 Ogawa ET, SmR SXTS, *lac-hlyA+/recA+* *wbeT^S158^* | ^1^ |
| *V. cholerae* Ogawa^V^ | HaitiV, O1 Ogawa ET, SmR SXTS, *lac-*Δ*hlyA/*Δ*recA wbeT^S158^* | This study |
| *V. cholerae* Inaba^V^ | HaitiV, O1 Inaba ET, SmR SXTS, *lac-* Δ*hlyA/*Δ*recA wbeT^S158P^* | This study |
| *V. cholerae* Hiko^V^ | HaitiV, O1 Hikojima ET, SmR SXTS, *lac-* Δ*hlyA/*Δ*recA wbeT^S158F^* | This study |
| *V. cholerae* N16961 | O1 Inaba ET, SmR | ^3^ |
| *V. cholerae* PIC018 | O1 Inaba ET | ^4^ |
| *V. cholerae* PIC158 | O1 Ogawa ET | ^4^ |

**Supplementary Table 2. Plasmids used in this study.** Cb = carbenicillin, Cm = chloramphenicol.

| **Plasmid** | **Notes** | **Source** |
| --- | --- | --- |
| pCVD442 | Allelic exchange vector for cloning *V. cholerae* mutant strains, CbR and *sacB+* | ^1^ |
| pCVD442_lacZ | Allelic exchange vector for in-frame deletion of *lacZ* | This study |
| pCVD442_hlyA | Allelic exchange vector for in-frame deletion of *hlyA* (VCA0219) | This study |
| pCVD442_recA | Allelic exchange vector for in-frame deletion of *recA* (VC0543) | This study |
| pCVD442_wbeT_S158P | Allelic exchange vector for replacement of Ogawa *wbeT* allele with Inaba *wbeT* allele; used for HaitiWT serotype cloning | This study |
| pCVD442_wbeT_S158F | Allelic exchange vector for replacement of Ogawa *wbeT* allele with Hikojima *wbeT* allele; used for HaitiWT serotype cloning | This study |
| pDS132 | Allelic exchange vector for cloning *V. cholerae* mutant strains, CmR and *sacB+* | ^5^ |
| pDS132_wbeT_S158P | Allelic exchange vector for replacement of Ogawa *wbeT* allele with Hikojima *wbeT* allele; used for HaitiV serotype cloning | This study |
| pDS132_wbeT_S158F | Allelic exchange vector for replacement of Ogawa *wbeT* allele with Hikojima *wbeT* allele; used for HaitiV serotype cloning | This study |
